# Drumming motor sequence training induces apparent myelin remodelling in Huntington’s disease: a longitudinal diffusion MRI and quantitative magnetization transfer study

**DOI:** 10.1101/2019.12.24.887406

**Authors:** Chiara Casella, Jose Bourbon-Teles, Sonya Bells, Elizabeth Coulthard, Greg D. Parker, Anne Rosser, Derek K. Jones, Claudia Metzler-Baddeley

**Affiliations:** Cardiff University Brain Research Imaging Centre (CUBRIC), School of Psychology, Cardiff University, Maindy Road, Cardiff, CF 24 4HQ, UK; The Hospital for Sick Children, Neurosciences and Mental Health, Toronto, M5G 1X8, Canada; Clinical Neurosciences, University of Bristol, Bristol, BS10 5NB, UK; School of Biosciences, Cardiff University, Museum Avenue, Cardiff, CF10 3AX, UK; Mary MacKillop Institute for Health Research, Australian Catholic University, Melbourne, Victoria 3065, Australia; Department of Neurology and Psychological Medicine, Hayden Ellis Building, Maindy Rd, CF24 4HQ

**Keywords:** Huntington’s disease, drumming training, white matter, myelin, diffusion MRI

## Abstract

1.

**Background:** Impaired myelination may contribute to Huntington’s disease (HD) pathogenesis. This study assessed differences in white matter (WM) microstructure between HD patients and controls, and tested whether drumming training stimulates WM remodelling in HD. Furthermore, it examined whether training-induced microstructural changes are related to improvements in motor and cognitive function.

**Methods:** Participants undertook two months of drumming exercises. Working memory and executive function were assessed before and after training. Changes in WM microstructure were investigated with diffusion tensor magnetic resonance imaging (DT-MRI)-based metrics, the restricted diffusion signal fraction (Fr) from the composite hindered and restricted model of diffusion (CHARMED) and the macromolecular proton fraction (MPF) from quantitative magnetization transfer (qMT) imaging. WM pathways linking the putamen and the supplementary motor area (SMA-Putamen), and three segments of the corpus callosum (CCI, CCII, CCIII) were studied using deterministic tractography. Baseline MPF differences between patients and controls were assessed with tract-based spatial statistics (TBSS).

**Results:** MPF was reduced in HD patients compared to controls in the mid-section of the CC in HD subjects at baseline, while a significantly greater change in MPF was detected in HD patients relative to controls in the CCII, CCIII, and the right SMA-putamen post-training. Further, although patients improved their drumming and executive function performance, such improvements did not correlate with microstructural changes.

**Conclusions:** Increased MPF suggests training-induced myelin changes in HD. Tailored behavioural stimulation may lead to neural benefits in early HD that could be exploited for delaying disease progression.

## 2. Introduction

### 2.1 The Pathology of Huntington’s disease

Huntington’s disease (HD) is a genetic, neurodegenerative disorder caused by an expansion of the CAG repeat within the *huntingtin* gene, leading to debilitating cognitive, psychiatric, and motor symptoms. In addition to striatal grey matter (GM) degeneration (Weaver et al., 2009), HD pathology has also been linked to white matter (WM) changes (Bardile et al., 2018; Bartzokis et al., 2007; Beglinger et al., 2007; Ciarmiello et al., 2006; Gregory et al., 2018; Paulsen et al., 2008; Rosas et al., 2018; Wang & Yang, 2019). WM is composed of axons and non-neuronal glia cells, such as myelin-producing oligodendrocytes, and it is unclear whether axons, myelin, or both are predominantly responsible for the WM abnormalities observed in HD (Gregory et al., 2018).

Some studies indicate that such changes are a result of Wallerian degeneration (Di Paola et al., 2014; Rosas et al., 2018; Weaver et al., 2009). However, an increasing body of research suggests that myelin-associated biological processes at the cellular and molecular level contribute to WM changes (Gómez-Tortosa et al., 2001; Huang et al., 2015; Jin et al., 2015; Myers et al., 1991; Simmons et al., 2007; Teo et al., 2016). Myelin is a multi-layered membrane sheath wrapping axons and is produced by oligodendrocytes. Axon myelination is vital during brain development and critical for healthy brain function, as it plays a fundamental role in the efficiency and speed of action potential propagation (Martenson, 1992). Oligodendrocyte/myelin dysfunction can slow down or stop otherwise fast axonal transport, which in turn can result in synaptic loss and eventually axonal degeneration (Han et al., 2010).

### 2.2 Interventions and Brain Plasticity

Currently, there are no disease-modifying treatments for HD. However, environmental stimulation and behavioural interventions may have the potential to reduce disease progression and delay disease onset (van Dellen et al., 2000; Yhnell et al., 2016). Converging evidence implicates myelin plasticity as one of the routes by which experience shapes brain structure and function (Caeyenberghs et al., 2018; Gibson et al., 2014; Lakhani et al., 2016; Mensch et al., 2015; Sampaio-Baptista et al., 2013; Scholz et al., 2009). Accordingly, plastic changes in myelination may be implicated in early adaptation and longer-term consolidation and improvement in motor tasks (Costa et al., 2004; Shmuelof & Krakauer, 2011; Steele et al., 2013; Yin et al., 2009). Changes in myelin-producing oligodendrocytes have been reported within the first hours of skill acquisition (Xiao et al., 2016); furthermore, alterations in grey and WM microstructure have been detected within 2 hours in people learning to play an action video game (Sagi et al., 2012), suggesting that experience can be quickly translated into adaptive changes in the brain.

This study assessed whether two months of drumming training could trigger WM microstructure changes, and potentially myelin remodelling, in individuals with HD. The present intervention was designed to exercise cognitive and motor functions including sequence and reversal learning, response speed and multi-tasking (Metzler-Baddeley et al., 2014), all of which rely on healthy functioning of cortico-basal ganglia loops that are known to be impaired in HD (Papoutsi et al., 2014). In brief, the intervention involves practising drumming patterns in ascending order of difficulty over a two-month period, and was previously found to induce WM changes in HD (Metzler-Baddeley et al. 2014). Based on reports of larger training-associated changes in structural MRI metrics in patient populations than in healthy subjects (Caeyenberghs et al., 2018), we hypothesised that changes in *microstructural* metrics would be more marked in patients than in healthy subjects.

Previous WM plasticity neuroimaging studies (Giacosa et al., 2016; Scholz et al., 2009) have predominantly employed indices from diffusion tensor MRI (DT-MRI) (Pierpaoli & Basser, 1996). However, while sensitive, such measures are not specific to changes in specific sub-compartments of WM microstructure, challenging the interpretation of any observed change in DT-MRI indices (De Santis et al., 2014; Wheeler-Kingshott & Cercignani, 2009). To improve compartmental specificity beyond DT-MRI, the present study explored changes in the macromolecular proton fraction (MPF) from quantitative magnetization transfer (qMT) (Sled, 2018) imaging and the restricted diffusion signal fraction (Fr) from the composite hindered and restricted model of diffusion (CHARMED) (Assaf & Basser, 2005). Fractional anisotropy (FA) and radial diffusivity (RD) from DT-MRI (Pierpaoli & Basser, 1996) were included for comparability with previous training studies (Lövdén et al., 2010; Scholz et al., 2009; Zatorre et al., 2012).

The MPF has been proposed as a proxy MRI marker of myelin (Serres et al., 2009). Accordingly, histology studies show that this measure reflects demyelination accurately in Shiverer moice (Ou et al., 2009), is sensitive to de-myelination in multiple sclerosis patients (Levesque et al., 2010) and reflects WM myelin content in post-mortem studies of multiple sclerosis brains (Schmierer et al., 2007). Fr, on the other hand, represents the fraction of signal-attenuation that can be attributed to restricted diffusion, which is presumed to be predominantly intra-axonal, and therefore provides a proxy measure of axonal density (Barazany et al., 2009).

Training effects were investigated in WM pathways linking the putamen and the supplementary motor area (SMA-Putamen), and within three segments of the corpus callosum (CCI, CCII, CCIII). The SMA has efferent and afferent projections to the primary motor cortex and is involved in movement execution, and previous work has reported altered DT-MRI metrics in the putamen-motor tracts of symptomatic HD patients (Poudel et al., 2014). The anterior and anterior-mid sections of the corpus callosum contain fibres connecting the motor, premotor and supplementary motor areas in each hemisphere (Hofer & Frahm, 2006). Previous work has demonstrated a thinning of the corpus callosum in post-mortem HD brains (Vonsattel & Difiglia, 1998), altered callosal DT-MRI metrics in both pre-symptomatic and symptomatic HD patients (Rosas et al., 2010; Phillips et al., 2013), and a correlation between these metrics and performance on motor function tests (Dumas et al., 2012). Given previous reports of an effect of motor learning on myelin plasticity (Lakhani et al., 2016;), we expected changes following training to be more marked in MPF, as compared to the other non-myelin sensitive metrics assessed in this study. We also investigated the relationship between training-associated changes in MRI measures, and changes in drumming performance and cognitive/executive function. Finally, as previous evidence has shown widespread reductions in MPF in premanifest and manifest HD patients (Bourbon-Teles et al., 2017), we used tract-based spatial statistics (TBSS) (Smith et al., 2006) to investigate patient-control differences in MPF before training, across the whole brain.

## 3. Materials and Methods

### 3.1 Participants

The study was approved by the local National Health Service (NHS) Research Ethics Committee (Wales REC 1 13/WA/0326) and all participants provided written informed consent. All subjects were drumming novices and none had taken part in our previously-reported pilot study (Metzler-Baddeley et al., 2014). Fifteen HD patients were recruited from HD clinics in Cardiff and Bristol. Genetic testing confirmed the presence of the mutant huntingtin allele. Thirteen age, sex, and education-matched healthy controls were recruited from the School of Psychology community panel at Cardiff University and from patients’ spouses, carers or family members. The inclusion criteria were the following: no history of head injury, stroke, cerebral haemorrhages or any other neurological condition; eligible for MRI scanning; stable medication for at least four weeks prior to the study.

Of the recruited sample, two patients were not MRI compatible, four withdrew during the study and one patient’s MRI data had to be excluded due to excessive motion. Therefore, while drumming performance and cognitive data from 11 patients were assessed, only 8 patients had a complete MRI dataset. One control participant was excluded due to an incidental MRI finding, two participants dropped out of the study and a fourth participant was not eligible for MRI. In total, we assessed drumming and cognitive tests performance in 8 controls, while MRI data from nine controls were available for analyses. Table 1 summarizes patients demographic and background clinical characteristics. Most patients were at early disease stages, however two were more advanced, as shown by their Total Motor Score (TMS; 69 and 40, respectively) and Functional Assessment Score (FAS; 18 and 17, respectively). Table 2 summarizes demographic variables and performance in the Montreal Cognitive Assessment (MoCA) (Nasreddine et al., 2005) and in the revised National Adult Reading Test (NART-R) (Nelson, 1991) for patients and controls. While the groups did not differ significantly in age, controls were on average slightly older, performed significantly better on the MoCA, and had a significantly higher NART-IQ than patients.

**Table 1.** *Demographics and background clinical information of the patients’ cohort*. Based on TMS and FAS, most of the patients were at early disease stages, however two of them were more advanced.

|  | Age | Length of CAG repeats | TMS | FAS |
| --- | --- | --- | --- | --- |
| Mean | 48.5 | 43.6 | 18.7 | 22.6 |
| (Range) | (22-68) | (40-51) | (0-69) | (17-25) |
| SD | 15.6 | 3.5 | 24.7 | 3.3 |

Abbreviations: CAG = cytosine-adenine-guanine; TMS = Total Motor Score out of 124 (the higher the scores the more impaired the performance); FAS = Functional Assessment Score out of 25 (the higher the scores the better the performance); SD = Standard Deviation.

**Table 2.** *Demographics and general cognitive profile of patients and controls. Both groups were matched for age, sex and years of education but the patient group had a lower* NART-IQ and *performed less well than the control group in the MoCA*.

| Mean<br>(SD, range) | Patients (n = 8) | Controls (n = 9) | Mann-Whitney U<br>(p-value) |
| --- | --- | --- | --- |
| Age | 48.5<br>(15.62, 22-68) | 52.6<br>(14.56, 22-68) | U = 31<br>(p = 0.673) |
| NART-IQ | 106.3<br>(13.13, 94-123) | 121.22<br>(4.32, 117-128) | U = 8<br>(p = 0.006) |
| MoCa | 23<br>(5.6, 14-29) | 27.67<br>(1, 26-29) | U = 14<br>(p = 0.036) |

Abbreviations: NART-IQ = verbal IQ estimate based on the National Adult Reading Test; MoCA = Montreal Cognitive Assessment score out of 30.

### 3.2 Training intervention: Drumming-based rhythm exercises

The rhythm exercise and drumming training described previously in Metzler-Baddeley et al. (2014) was applied. Participants were provided with twenty-two 15 min training sessions on CDs, a pair of Bongo drums and a drumming diary and could practise at home. They were asked to exercise for 15 min per day, 5 times per week, for 2 months and to record the date and time of each exercise in their diary. Each training session introduced a drumming pattern based on one of the following rhythms: Brazilian samba, Spanish rumba, West-African kuku and Cuban son. After a brief warm up, trainees were encouraged to drum along with the instructor, initially with each hand separately and then with both hands alternating, starting with the dominant hand first and then reversing the order of the hands. The first exercises were based on very simple, slow, and regular patterns but the level of complexity and speed increased over the training sessions. Importantly, each individual progressed through the training adaptively at their own pace i.e., as long as they exercised for the specified time, they could repeat each session as often as they felt necessary to master it. To maintain engagement and motivation, the training incorporated pieces of music based on rhythms participants had learned and could drum along to. The researcher (JBT) supervised the first training sessions and then remained in regular telephone contact (at least once a week) with each participant throughout the intervention. Whenever possible, carers and/or spouses were involved in the study to support the training. Control participants started with Session 3 since the first two exercises were built on a very low level of complexity, with slow, regular patterns of movement required, and were therefore designed for patients.

### 3.3 Drumming assessment

Progress in drumming ability was assessed by digitally recording participants’ drumming performance for three patterns of ascending levels of difficulty (easy, medium and hard), which were not part of the training sessions, at baseline and after the training. Each recording was judged by an independent rater, blind to group and time, according to an adopted version of the Trinity College London marking criteria for percussion (2016) (www.trinitycollege.com; see Supplementary Figure 1 for details).

### 3.4 Cognitive assessments

Different aspects of cognition and executive function were assessed before and after the training as described in Metzler-Baddeley et al. (2014). Multi-tasking was assessed with a dual task requiring simultaneous box crossing and digit sequences repetition (Baddeley, 1996). Attention switching was assessed with the trails test (VT) requiring the verbal generation of letter and digit sequences in alternate order relative to a baseline condition of generating letter or digit sequences only (Baddeley, 1996). Distractor suppression was tested with the Stroop task involving the naming of incongruent ink colours of colour words (Trenerry et al., 1989). Verbal and category fluency were tested using the letter cues “F”, “A”, “S” and “M”, “C”, “R” as well as the categories of “animals” and “boys’ names” and “supermarket items” and “girls’ names” respectively (Baldo et al., 2001). In total, we assessed 7 outcome variables, and percentage change scores in performance were computed for each of these variables (Table 3).

**Table 3.**
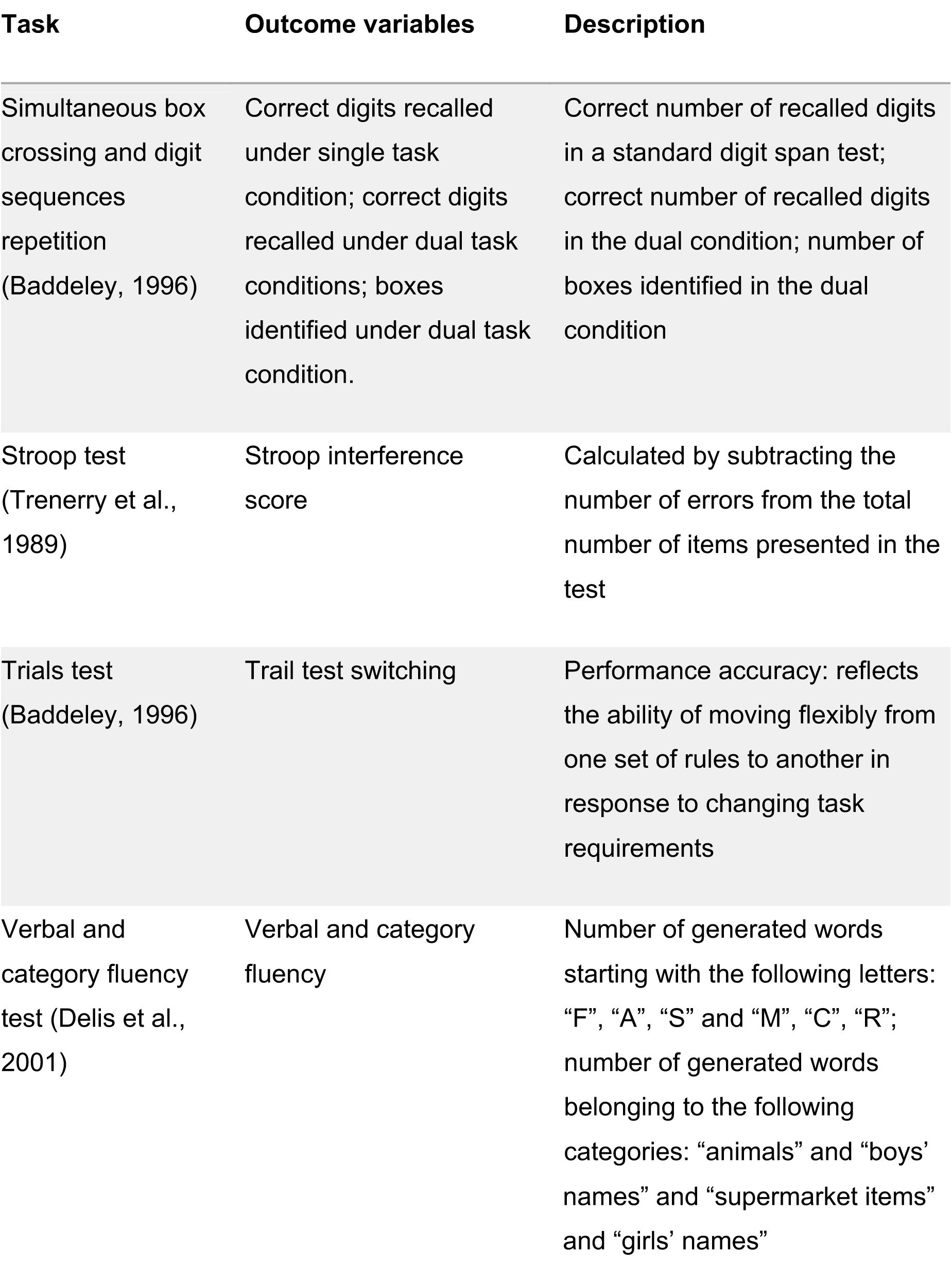
Cognitive outcome variables assessed in this study. Tests were carried out before and after the training, and a percentage change score was computed for each variable.

### 3.5 MRI data acquisition

MRI data were acquired on a 3 Tesla General Electric HDx MRI system (GE Medical Systems, Milwaukee) using an eight channel receive-only head RF coil at the Cardiff University Brain Research Imaging Centre (CUBRIC). The MRI protocol comprised the following images sequences: a high-resolution fast spoiled gradient echo (FSPGR) T_1_-weighted (T_1_-w) sequence for registration; a diffusion-weighted spin-echo echo-planar sequence (SE\EPI) with 60 uniformly distributed directions (b?=?1200 s/mm^2^), according to an optimized gradient vector scheme (Jones et al., 1999); a CHARMED acquisition with 45 gradient orientations distributed on 8 shells (maximum b-value = 8700s/mm^2^) (Assaf and Basser, 2005); and a 3D MT-weighted fast spoiled gradient recalled-echo (FSPGR) sequence (Cercignani and Alexander, 2006). The acquisition parameters of all scan sequences are reported in Table 4. Diffusion data acquisition was peripherally gated to the cardiac cycle. The off-resonance irradiation frequencies (Θ) and their corresponding saturation pulse amplitude (ΔSAT) for the 11 Magnetization transfer (MT) weighted images were optimized using Cramer-Rao lower bound optimization (Cercignani & Alexander, 2006).

**Table 4.** Scan parameters. All sequences were acquired at 3T. For each of the sequences, the main acquisition parameters are provided. T_1_-w: T_1_-weighted; MT-w: MT-weighted; FSPGR: fast spoiled gradient echo; SE: spin-echo; EPI: echo-planar imaging; SPGR: spoiled gradient recalled-echo; FoV: field of view; TE: echo time; TR: repetition time.

|  | T<sub>1</sub>-w | DTI | CHARME<br>D | T1 map | MT-w | B0 map |
| --- | --- | --- | --- | --- | --- | --- |
| Pulse<br>sequence | FSPGR | SE\EPI | SE\EPI | SPGR<br>(3D) | FSPGR<br>(3D) | SPGR<br>(3D) |
| Matrix<br>size | 256×256 | 96×96 | 96×96 | 96×96×60 | 96×96×60 | 128×128 |
| FoV<br>(mm) | 230 | 230 | 230 | 240 | 240 | 220 |
| Slices | 172 | 60 | 60 | - | - | - |
| Slice<br>thickness<br>(mm) | 1 | 2.4 | 2.4 | - | - | - |
| TE,TR<br>(ms) | 7.8, 2.9 | 87,<br>16000 | 126, 17000 | 6.85, 1.2 | 2.18,25.82 | TE: 9 & 7<br>TR: 20 |
| **Off-resonance pulses (Hz/°)** | - | - | - | - | 1000/332,<br>1000/333,<br>12062/628,<br>47185/628,<br>56363/332,<br>2751/628,<br>1000/628,<br>1000/628,<br>2768/628,<br>2791/628,<br>2887/628 | - |
| **Flip angles (°)** | 20 | 90 | 90 | 15,7,3 | 5 | 90 |

### 3.6 MRI data processing

The diffusion-weighted data were corrected for distortions induced by the diffusion-weighted gradients, artefacts due to head motion and EPI-induced geometrical distortions by registering each image volume to the T_1_-w anatomical images (Irfanoglu et al., 2012), with appropriate reorientation of the encoding vectors (Alexander Leemans & Jones, 2009), all done in ExploreDTI (Version 4.8.3) (Leemans, Jeurissen, Sijbers, & Jones, 2009). A two-compartment model was fitted to derive maps of FA and RD in each voxel (Pasternak et al., 2009). CHARMED data were corrected for motion and distortion artefacts according to the extrapolation method of Ben-Amitay, Jones, and Assaf (2012). The number of distinct fiber populations (1, 2, or 3) in each voxel was obtained using a model selection approach (De Santis et al., 2014) and Fr was calculated per voxel with an in-house software (De Santis et al., 2014) coded in MATLAB (The MathWorks, Natick, MA). MT-weighted SPGR volumes for each participant were co-registered to the MT-volume with the most contrast using an affine (12 degrees of freedom, mutual information) registration to correct for inter-scan motion using Elastix (Klein et al., 2010). The 11 MT-weighted SPGR images and T_1_ map were modelled by the two pool Ramani’s pulsed MT approximation (Henkelman et al., 1993; Ramani et al., 2002), which included corrections for amplitude of B_0_ field inhomogeneities. This approximation provided MPF maps, which were nonlinearly warped to the T_1_-w images using the MT-volume with the most contrast as a reference using Elastix (normalized mutual information cost function) (Klein et al., 2010).

### 3.7 Deterministic Tractography

Training-related changes in FA, RD, Fr, and MPF were quantified using a tractography approach in pathways interconnecting the putamen and the supplementary motor area bilaterally (SMA-Putamen), and within three segments of the corpus callosum (CCI, CCII and CCIII) (Hofer & Frahm, 2006) (Figure 1).

**Figure 1.**
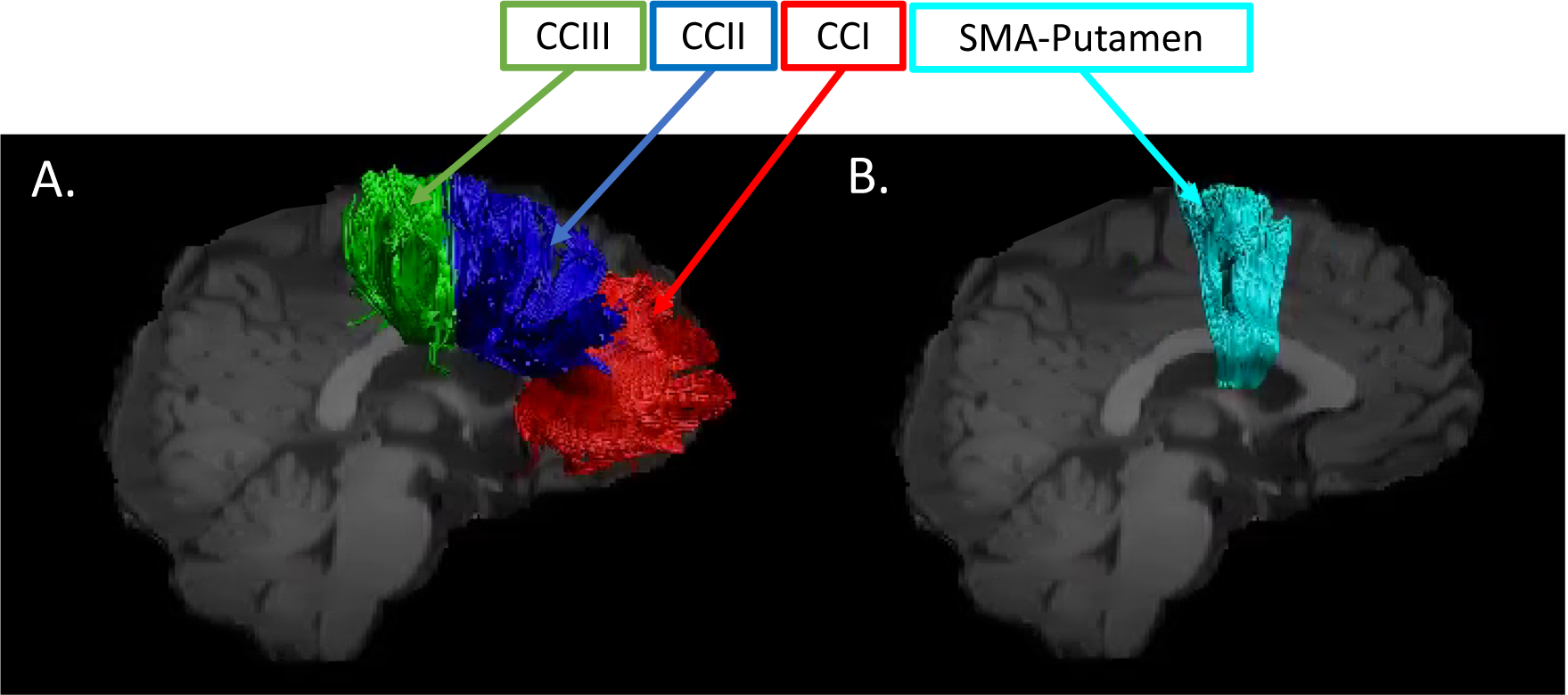
White matter pathway regions of interest. Sagittal views of the reconstructed WM pathways displayed on a T_1_-weighted image for one control participant. (A) CCI, CCII, and CCIII (Hofer and Frahm, 2006): CCI is the most anterior portion of the CC and maintains prefrontal connections between both hemispheres; CCII is the portion that maintains connections between premotor and supplementary motor areas of both hemispheres. CCIII maintains connections between primary motor cortices of both hemispheres. (B) SMA-putamen pathway: this pathway has efferent and afferent projections to the primary motor cortex and is involved in movement execution.

Whole brain tractography was performed for each participant in their native space using the damped Richardson-Lucy algorithm (Dell’acqua et al., 2010), which allows the recovery of multiple fiber orientations within each voxel including those affected by partial volume. The tracking algorithm estimated peaks in the fiber orientation density function (fODF) by selecting seed points at the vertices of a 2×2×2 mm grid superimposed over the image and propagated in 0.5-mm steps along these axes re-estimating the fODF peaks at each new location (Jeurissen et al., 2011). Tracks were terminated if the fODF threshold fell below 0.05 or the direction of pathways changed through an angle greater than 45° between successive 0.5 mm steps. This procedure was then repeated by tracking in the opposite direction from the initial seed-points.

The WM tracts of interest were extracted from the whole-brain tractograms by applying way-point regions of interest (ROI) (Catani et al., 2002). These were drawn manually by one operator (JBT) blind to the identity of each dataset on color-coded fiber orientation maps in native space guided by the following anatomical landmark protocols (Figure 2).

**Figure 2.**
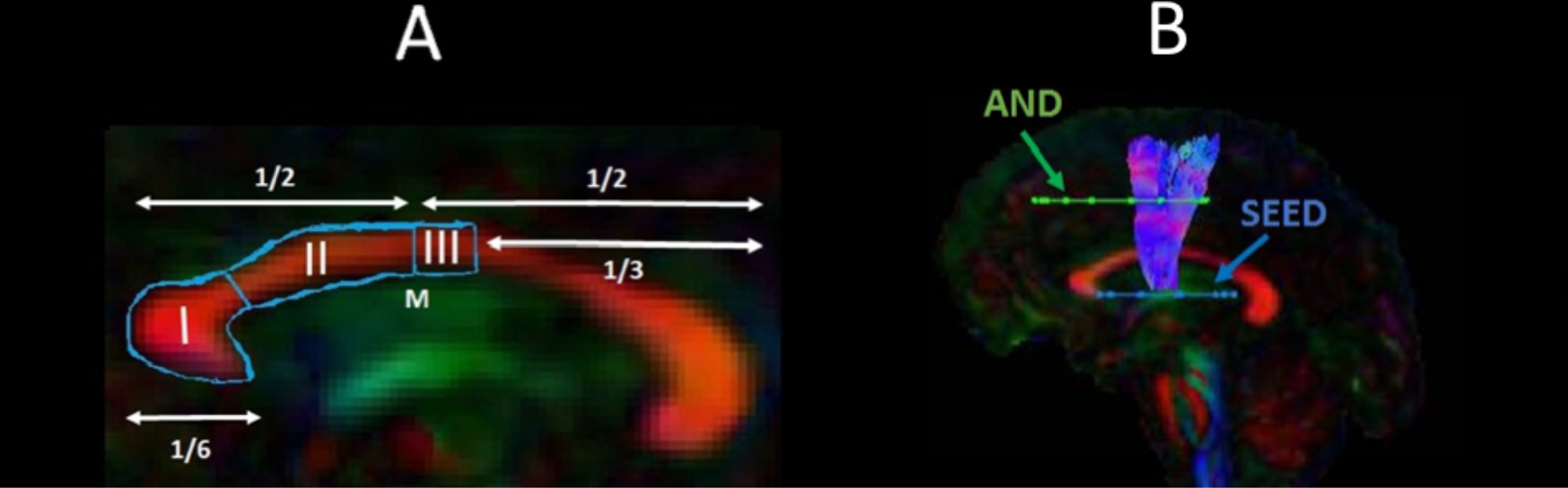
Sagittal views of the tractography protocols. (A) CCI, CCII and CCIII (B) SMA - putamen pathway. Booleian logic OR waypoint regions of interest gates are illustrated in blue; AND gates in green. M = Midline.

#### 3.7.1 Corpus callosum

Reconstruction of the CC segments followed the protocol of Hofer and Frahm (Hofer & Frahm, 2006) as illustrated in Figure 2A. Segment reconstructions were visually inspected and, if necessary, additional gates were placed to exclude streamlines inconsistent with the known anatomy of the CC.

#### 3.7.2 SMA-putamen pathway

One axial way-point ROI was placed around the putamen and one axial ROI around the supplementary motor cortex (Leh et al., 2007) (Figure 2B). A way-point gate to exclude fibres projecting to the brain stem was placed inferior to the putamen.

### 3.8 Statistical analyses

Statistical analyses were carried out in R Statistical Software (Foundation for Statistical Computing, Vienna, Austria).

#### 3.8.1 Assessment of training effects on drumming performance

Improvements in drumming performance were analysed with a two-way mixed analysis of variance (ANOVA) testing for the effects of group (HD/controls), time of assessment (before/after the training) and group by time interaction effects. Furthermore, we compared our results to the ones obtained by running a robust mixed ANOVA, using the bwtrim R function from the WRS2 package (Mair & Wilcox, 2020). This implements robust methods for statistical estimation and therefore provides a good option to deal with data presenting small sample sizes, skewed distributions and outliers (Wilcox, 2011). Significant effects were further explored with post-hoc paired and independent t-tests. The reliability of the post-hoc analyses was assessed with bootstrap analysis based on 1000 samples and the 95% confidence interval (CI) of the mean difference is provided for each significant comparison.

#### 3.8.2 Assessment of group differences in the effect of training on cognitive performance

Performance measures in executive function tasks have been shown to share underlying cognitive structures (Testa et al., 2012). Therefore, PCA was employed to reduce the complexity of the cognitive data and hence the problem of multiple comparisons as well as to increase experimental power. PCA was run on change scores for all participants across both groups. Due to the relatively small sample size, we first confirmed with the Kaiser-Meyer-Olkin (KMO) test that our data was suited for PCA. Subsequently, we followed guidelines to limit the number of extracted components (Preacher & MacCallum, 2002; Winter, Dodou, & Wieringa, 2009), as follows: first, we employed the Kaiser criterion of including all components with an eigenvalue greater than 1; second, we inspected the Cattell scree plot (Cattell, 1966) to identify the minimal number of components that accounted for most variability in the data; third, we assessed each component’s interpretability. A PCA procedure with orthogonal Varimax rotation of the component matrix was used. Loadings that exceeded a value of 0.5 were considered as significant.

Next, we assessed group differences in the component scores with permutation analyses, to understand whether the training had differentially affected HD patients as compared to controls. Significant group differences were tested using 5,000 permutations. Permutation testing relies only on minimal assumptions and can therefore be applied when the assumptions of a parametric approach are untenable such as in the case of small sample sizes. Multiple comparison correction was based on a 5% false discovery rate (FDR) using the Benjamini-Hochberg procedure (Benjamini & Hochberg, 1995).

#### 3.8.3 Training effects on WM microstructure

Median measures of FA, RD, Fr and MPF were derived for each of the reconstructed tracts in ExploreDTI (Leemans, Jeurissen, Sijbers, & Jones, 2009). A percentage change score in these measures between baseline and post-training was calculated in each tract (CCI, CCII, CCIII, left and right SMA-Putamen).

Previous research has shown that variation in the microstructural properties of WM may represent a global effect, rather than being specific to individual tracts, and that WM measures are highly correlated across WM areas (Lövdén et al., 2010; Penke et al., 2010; Wahl et al., 2010). Therefore, we inspected the inter-tract correlation for each of microstructural metric and found that MPF values were highly correlated, whereas this was not true for the other metrics (Figure 3). Hence, percentage change scores in MPF across the different tracts were transformed with PCA to extract meaningful anatomical properties, following the procedure described above for the PCA of cognitive change scores. PC scores for each participant were then used as dependent variables in a permutation-based analysis using 5,000 permutations to assess group differences in training associated changes in MPF. Finally, as a post-hoc exploration, we looked for between-groups differences in MPF changes in the individual tracts using 5000 permutations.

**Figure 3.**
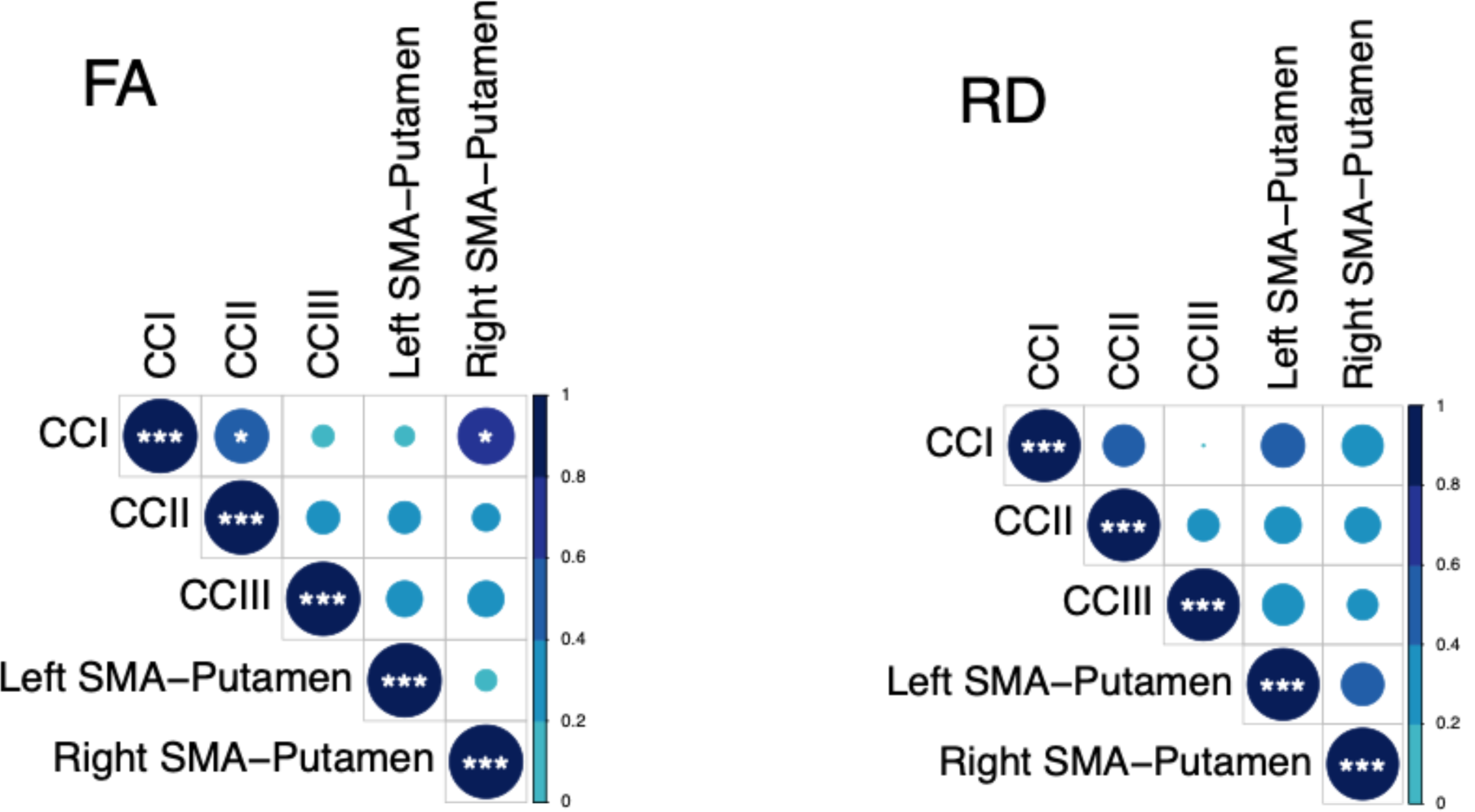
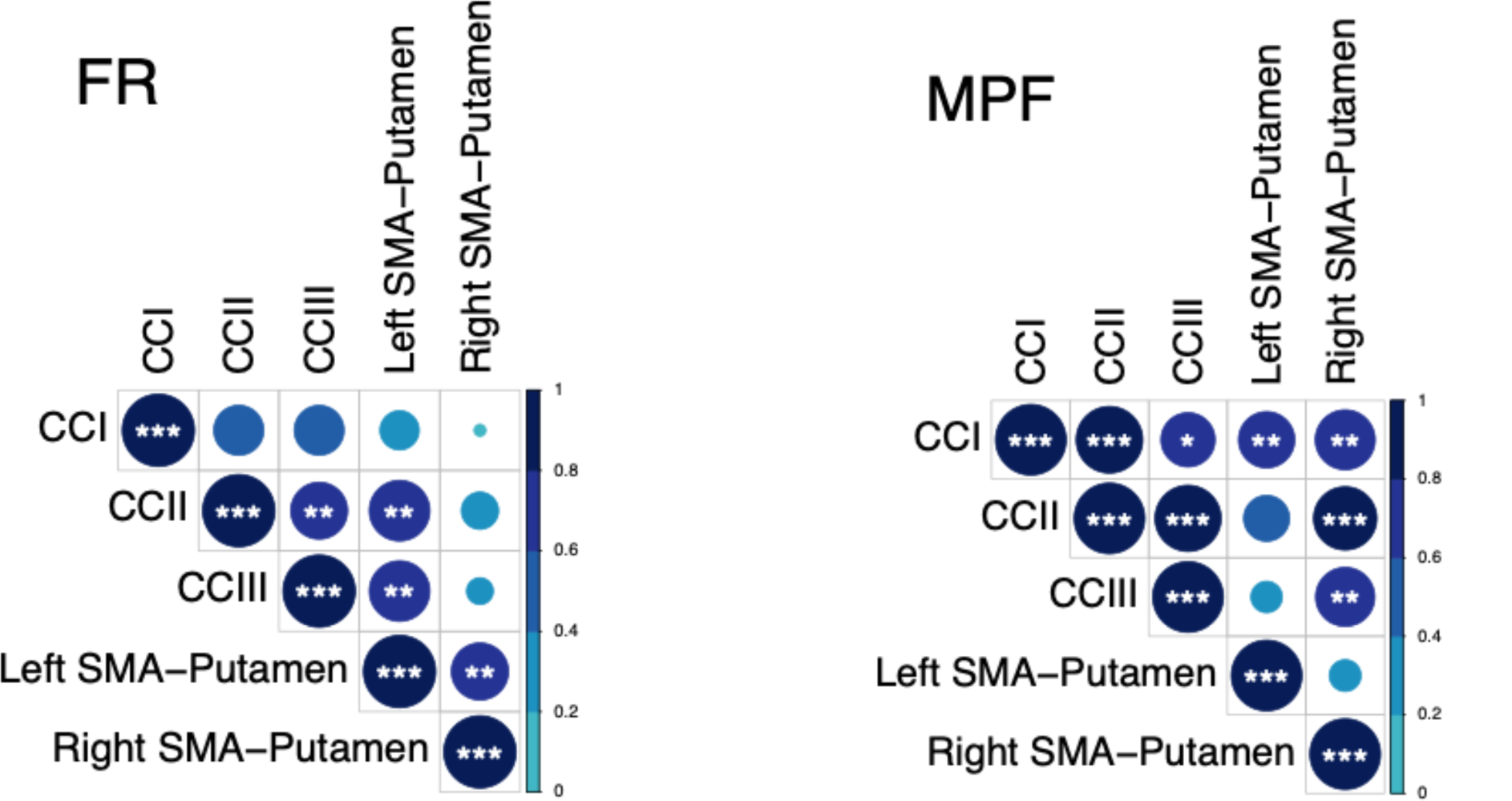
Correlation matrices for the MRI metrics investigated across the different WM pathways. Colour intensity and the size of the circles are proportional to the strength of the correlation. * p < 0.05, ** p < 0.01, *** p < 0.001. The absolute correlation coefficient is plotted. MPF values were highly correlated across tracts, whereas this was not true for the other metrics

Training-associated changes in FA, Fr and RD were investigated with permutation analyses separately for each tract. Significant group differences in these measures were tested using 5,000 permutations. Multiple comparison correction was based on a 5% FDR using the Benjamini-Hochberg procedure (Benjamini & Hochberg, 1995).

TBSS (Smith et al., 2006) was carried out to investigate baseline differences in MPF between HD subjects and healthy controls. To produce significance maps, a voxel-wise analysis was performed on the MPF projected 4D data for all voxels with FA ≥ 0.20 to exclude peripheral tracts where significant inter-subject variability exists. Inference based on permutations (5,000 permutations) and threshold-free-cluster-enhancement was used. The significance level was set at p < 0.05 and corrected by multiple comparisons (family-wise error, FWE).

#### 3.8.4 Relationship between changes in MRI measures and changes in drumming and cognitive performance

We computed percentage change scores for the drumming performance, in the same way cognitive change scores were calculated. Scores were computed for the easy test pattern in patients and for the medium test pattern in controls, as these training patterns showed a significant improvement in the two groups, respectively. Spearman correlation coefficients were calculated between drumming and cognitive performance, and microstructural components that showed significant group differences, to assess whether microstructural changes were related to any drumming and/or cognitive benefits of the training.

## 4. Results

### 4.1 Training effects on drumming performance

The mixed ANOVA of drumming performance for the easy and medium test pattern showed a significant effect of group [easy: F(1,17) = 22.3, p < 0.001; medium: F(1,17) = 13.1, p = 0.002] and time [easy: F(1,17) = 12.83, p = 0.004; medium: F(1,17) = 13.4, p = 0.002] but no interaction (easy: p = 0.8; medium: p = 0.3). For the hard test pattern there was only a significant effect of group [F(1,17) = 9.95, p = 0.006] but not of time (p = 0.1) and there was no interaction (p = 0.4). Results from the robust mixed ANOVA were largely consistent with the above. Specifically, the easy and medium test patterns showed a significant effect of group (easy: p = 0.002; medium: p = 0.02) and time (easy: p = 0.04; medium: p = 0.049) but no interaction (easy: p = 0.45; medium: p = 0.69). The hard test pattern showed a significant effect of group (p = 0.02) but not of time (p = 0.22) and no interaction (p = 0.8). Figure 4 summarises the average drumming performance per group and time point. Overall patients’ drumming performance was poorer than controls. Patients improved their drumming performance significantly for the easy pattern [t(10) = 2.7, p = 0.02; 95% CI of mean difference: 1.5 – 7.8] and controls for the medium pattern [t(7) = 3.8, p = 0.01; 95% CI of mean difference: 2.8 – 8.5].

**Figure 4.**
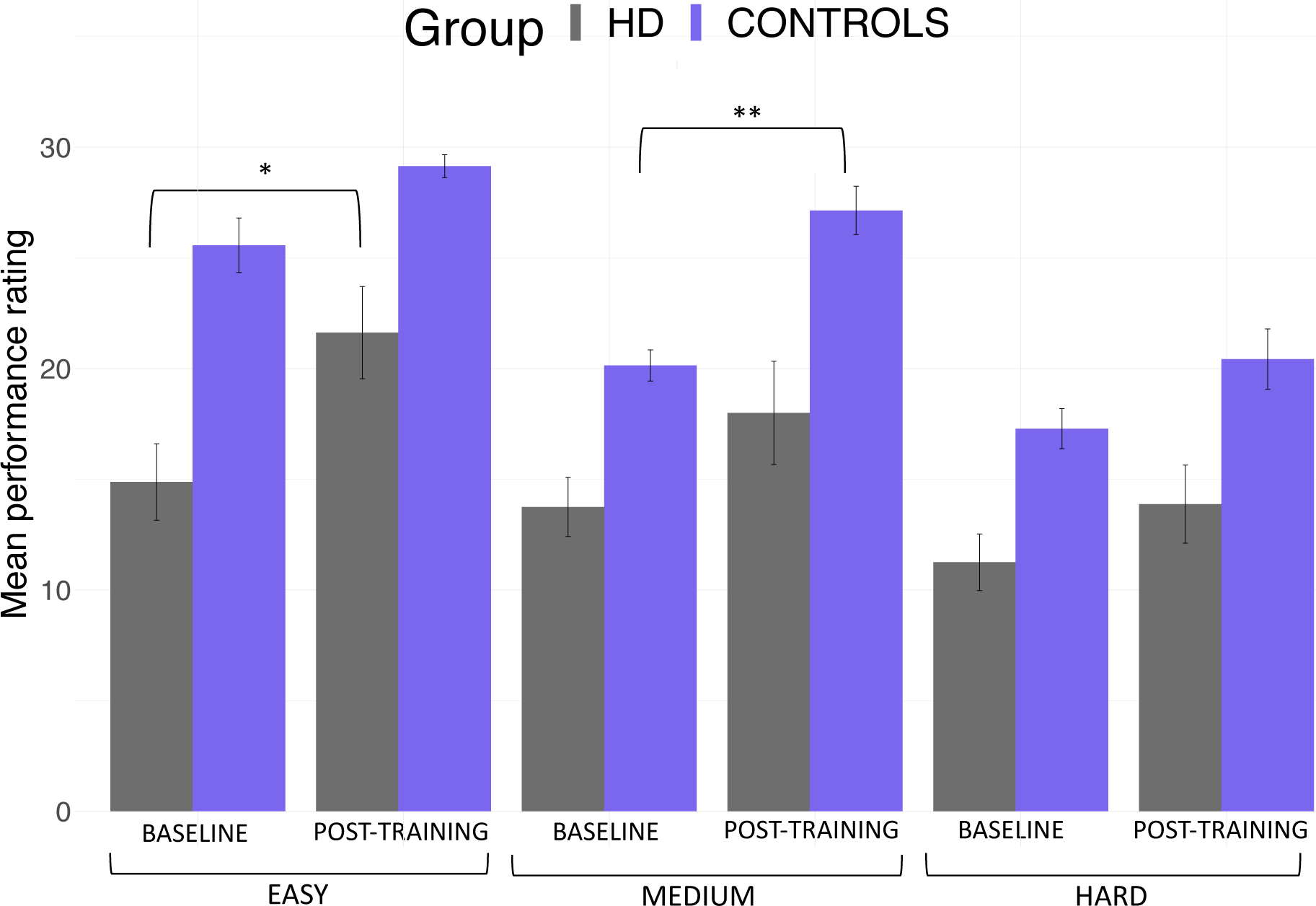
Mean ratings for drumming performance according to the Trinity College London marking criteria for percussion (2016) as a function of group and time point. Patients improved their drumming performance significantly for the easy test pattern and controls for the medium difficult test pattern. * p < 0.05, ** p < 0.01, bootstrapping based on 1000 samples.

### 4.2 Group differences in the effect of training on cognitive performance

Three components, accounting for 79% of the variance in performance improvement in the cognitive benchmark tests, were extracted. The first component loaded highly on performance changes in the dual task (total number of boxes identified under dual task condition), the Stroop task (Stroop interference score), and the trails making task (Trail test switching). Since these variables all measure executive functions including focused attention and distractor suppression, the first component was labelled “executive” component. The second component loaded on variables reflecting the ability to correctly recall digits sequences (i.e. number of correct digits recalled under single and dual task condition) and was therefore labelled “working memory capacity” component. Finally, the third extracted component loaded highly on verbal and category fluency and was therefore labelled “fluency” component (Table 5).

**Table 5.** Rotated Component Loadings on Change in the Cognitive Benchmark Tests. Significant loadings (>0.5) are highlighted in bold.

| % Change | Executive | Working memory capacity | Fluency |
| --- | --- | --- | --- |
| Total box (dual) | **0.864** | 0.022 | 0.419 |
| Stroop interference score | **0.811** | -0.270 | -0.267 |
| Trail test switching | **0.731** | -0.470 | 0.162 |
| Correct digits<br>under single task<br>condition | 0.201 | **0.904** | 0.129 |
| Correct digits<br>under dual task<br>condition | -0.193 | **0.855** | -0.018 |
| Category fluency | -0.070 | -0.138 | **0.817** |
| Verbal fluency | -0.026 | -0.232 | **-0.799** |

We tested whether the two groups differed in terms of post-training cognition changes, by running permutation analyses on the individual scores for the three extracted components. The two groups differed in the executive component, t = -1.03, p = 0.008, FDR-corrected p = 0.024. However, no significant group differences were detected in the other two components (Working Memory capacity: t = -0.22, p = 0.3296, FDR-corrected p = 0.3296; Fluency: t = -0.39, p = 0.242 FDR corrected p = 0.3296).

### 4.3 Training effects on WM microstructure

Table 6 reports a summary of the training associated changes in FA, RD, Fr and MPF, across the different tracts.

**Table 6.**
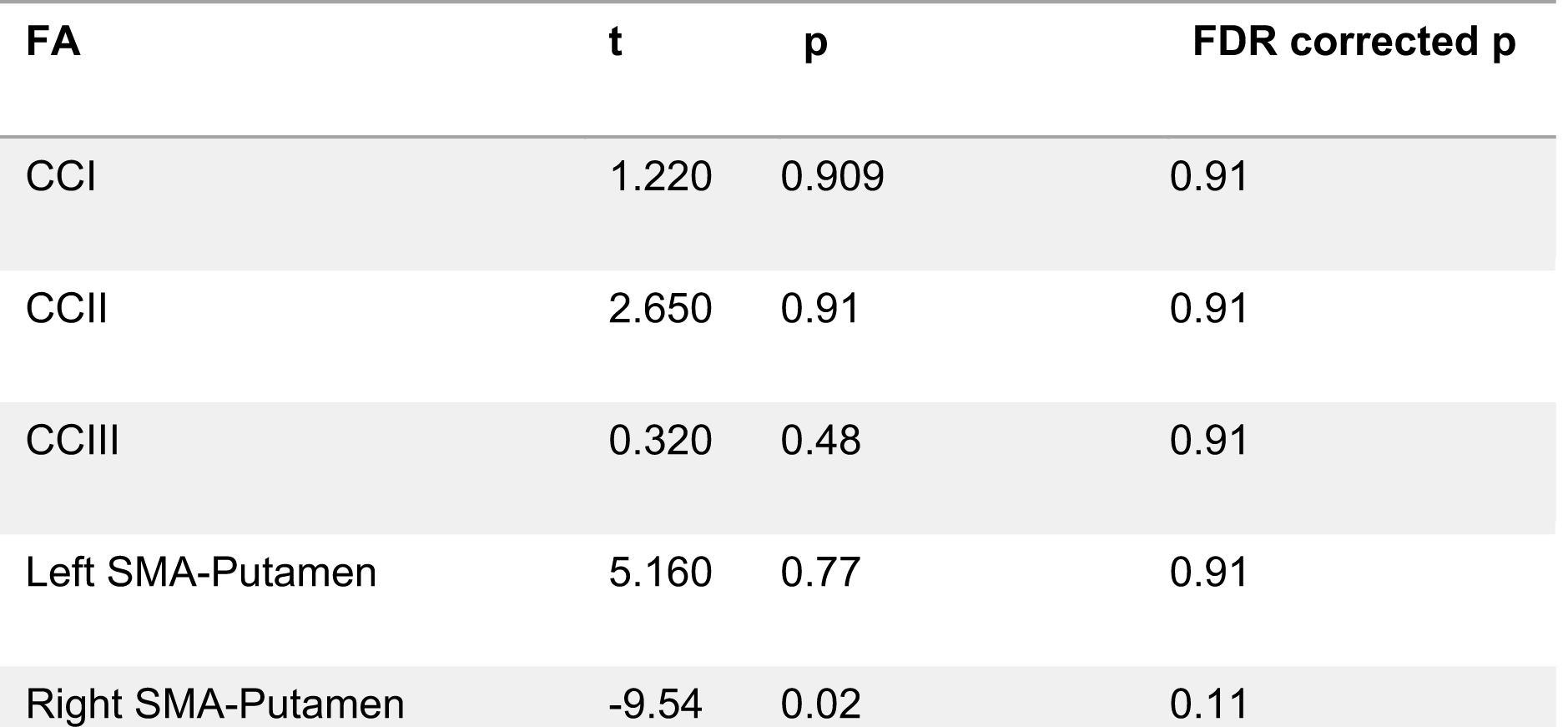
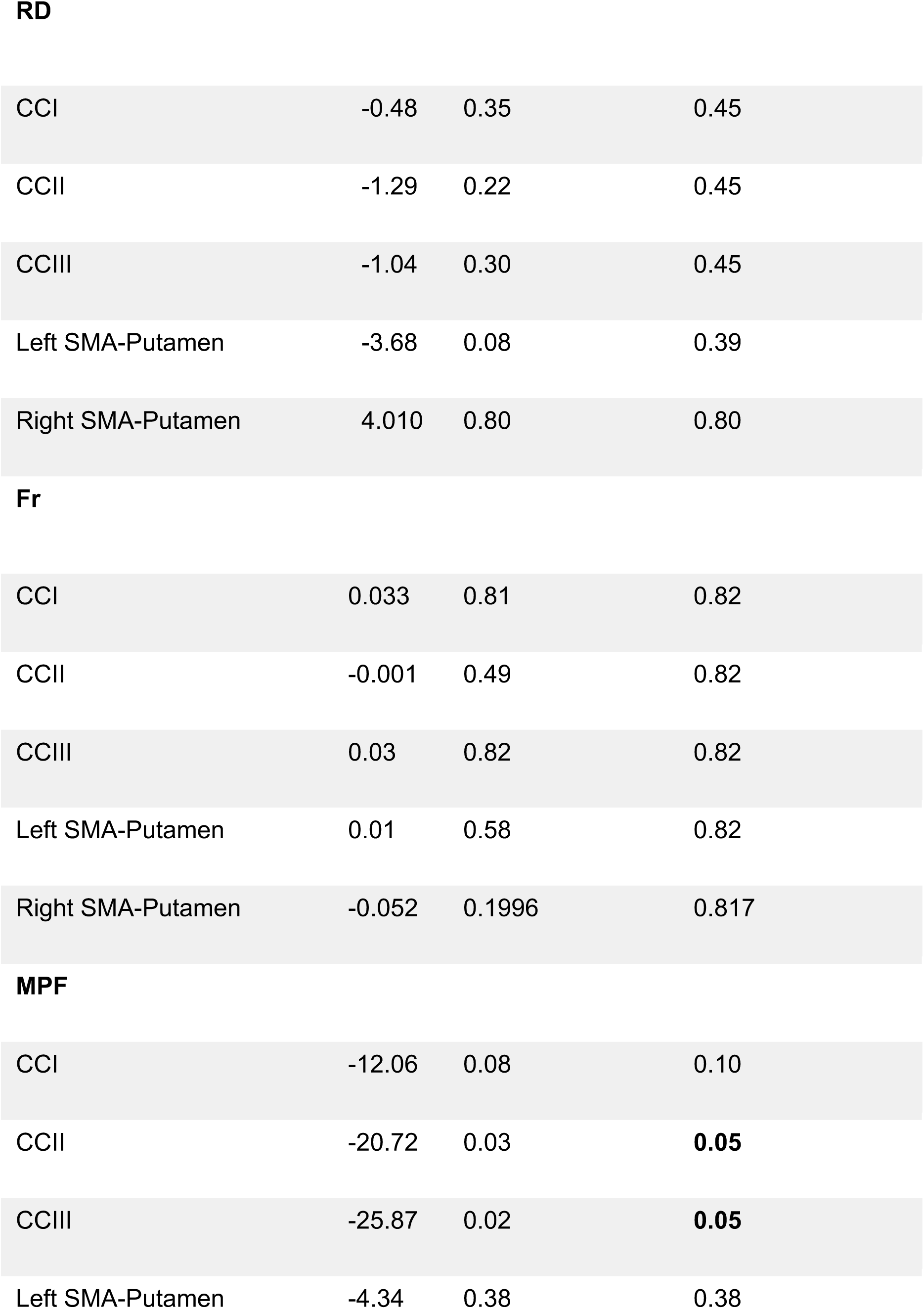
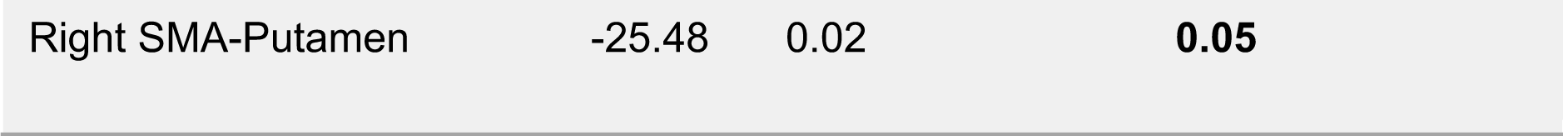
Summary statistics for the permutation analysis of training effects on FA, RD, Fr and MPF, across the investigated tracts.

| FA | t | p | FDR corrected p |
| --- | --- | --- | --- |
| CCI | 1.220 | 0.909 | 0.91 |
| CCII | 2.650 | 0.91 | 0.91 |
| CCIII | 0.320 | 0.48 | 0.91 |
| Left SMA-Putamen | 5.160 | 0.77 | 0.91 |
| Right SMA-Putamen | -9.54 | 0.02 | 0.11 |
| **RD** |  |  |  |
| CCI | -0.48 | 0.35 | 0.45 |
| CCII | -1.29 | 0.22 | 0.45 |
| CCIII | -1.04 | 0.30 | 0.45 |
| Left SMA-Putamen | -3.68 | 0.08 | 0.39 |
| Right SMA-Putamen | 4.010 | 0.80 | 0.80 |
| **Fr** |  |  |  |
| CCI | 0.033 | 0.81 | 0.82 |
| CCII | -0.001 | 0.49 | 0.82 |
| CCIII | 0.03 | 0.82 | 0.82 |
| Left SMA-Putamen | 0.01 | 0.58 | 0.82 |
| Right SMA-Putamen | -0.052 | 0.1996 | 0.817 |
| **MPF** |  |  |  |
| CCI | -12.06 | 0.08 | 0.10 |
| CCII | -20.72 | 0.03 | **0.05** |
| CCIII | -25.87 | 0.02 | **0.05** |
| Left SMA-Putamen | -4.34 | 0.38 | 0.38 |
| Right SMA-Putamen | -25.48 | 0.02 | **0.05** |

#### 4.3.1 Training-associated group differences in FA

Permutation analyses of FA changes across the different tracts revealed no significant differences between HD and control groups [CCI: t = 1.22, p = 0.91 (FDR-corrected); CCII: t = 2.65, p = 0.91 (FDR-corrected); CCIII: t = 0.325, p = 0.13 (FDR-corrected); right SMA-Putamen: t = -9.54, p = 0.10 (FDR-corrected); left SMA-Putamen: t = 5.16, p = 0.77 (FDR-corrected).

#### 4.3.2 Training-associated group differences in RD

There were no significant differences in RD changes following training between HD patients and controls [CCI: t = -0.48, p = 0.45 (FDR-corrected); CCII: t = -1.29, p = 0.45 (FDM-corrected); CCIII: t = -1.04, p = 0.45 (FDR-corrected); right SMA-Putamen, t = 4.01, p = 0.81 (FDR-corrected); left SMA-Putamen, t = -3.68, p = 0.39 (FDR-corrected).

#### 4.3.3 Training-associated group differences in Fr

Permutation analyses of Fr changes across the different tracts revealed no significant differences between HD and control groups [CCI: t = 3.39, p = 0.82 (FDR-corrected; CCII: t = -0.17, p = 0.82 FDR-corrected; CCIII: t = 3.08, p = 0.82 (FDR-corrected); right SMA-Putamen: t = -5.24, p = 0.82 (FDR-corrected); left SMA-Putamen: t = 1.05, p = 0.82 (FDR-corrected)].

#### 4.3.4 Training-associated group differences in MPF

PCA of change scores in MPF revealed one single component explaining 70.2% of the variance. This component presented high loadings from all the tracts investigated. A significant group difference was found for the MPF change-score component, indicating that HD patients presented significantly greater MPF changes in response to training, as compared to controls [t(14) = -1.743, n = 17, p = 0.03].

Finally, we found a significant difference in mean MPF change scores between the two groups for CCII [t(14) = -20.72, p=0.04], CCIII [t(14) = -25.87, p=0.04], and the right SMA-putamen pathway [t(14) = -25.48, p=0.04] after FDR correction, therefore indicating that there was a differential group effect of training on MPF within these tracts (Figure 5).

**Figure 5.**
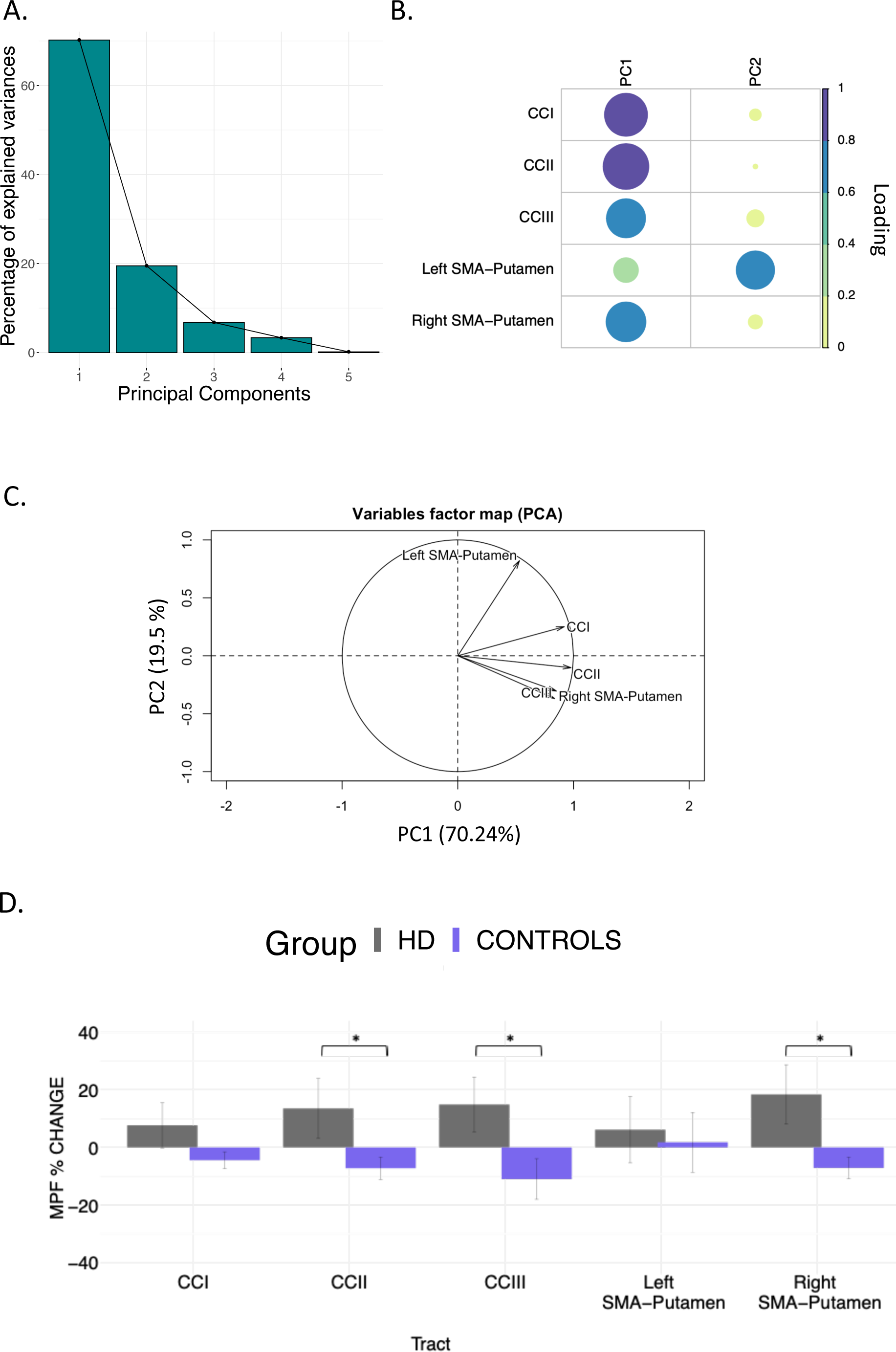
MPF changes scores: PCA scree plot (A); plot summarising how each variable is accounted for in every principal component - colour intensity and the size of the circles are proportional to the loading: PC1 loads on CCI, CCII, CCIII and right SMA-Putamen, while PC2 loads mostly on the left SMA-Putamen; the absolute correlation coefficient is plotted (B); correlation circle, interpreted as follows: 1) positively correlated variables are grouped together, 2) negatively correlated variables are positioned on opposite sides of the plot origin (opposite quadrants), 3) the distance between variables and the origin measures the quality of the variable on the factor map. Variables that are away from the origin are well represented on the factor map (C); Bar graph of the percentage change in MPF across the inspected tracts; Error bars represent the standard error; training was associated with a significantly greater change in MPF in CCII, CCIII, and right SMA-Putamen; * (p<0.05), results corrected for multiple comparisons with FDR (D).

### 4.4 Relationship between training-associated changes in MRI measures, and changes in drumming and cognitive performance

We did not find a significant association between the ‘MPF’ component scores and improvement in drumming performance (PC1: rho = -0.14, p > 0.05). Moreover, although no significant correlation was observed between the ‘Executive’ and the ‘MPF’ component scores, there was a positive trend (rho = .348, p = .171)

### 4.5 Baseline differences in MPF

Baseline MPF was reduced in the HD group compared to controls, in the midbody of the CC (t = 3.13, p = .05, FWE corrected). Figure 6 shows the areas with reduced MPF in HD patients, in blue.

**Figure 6.**
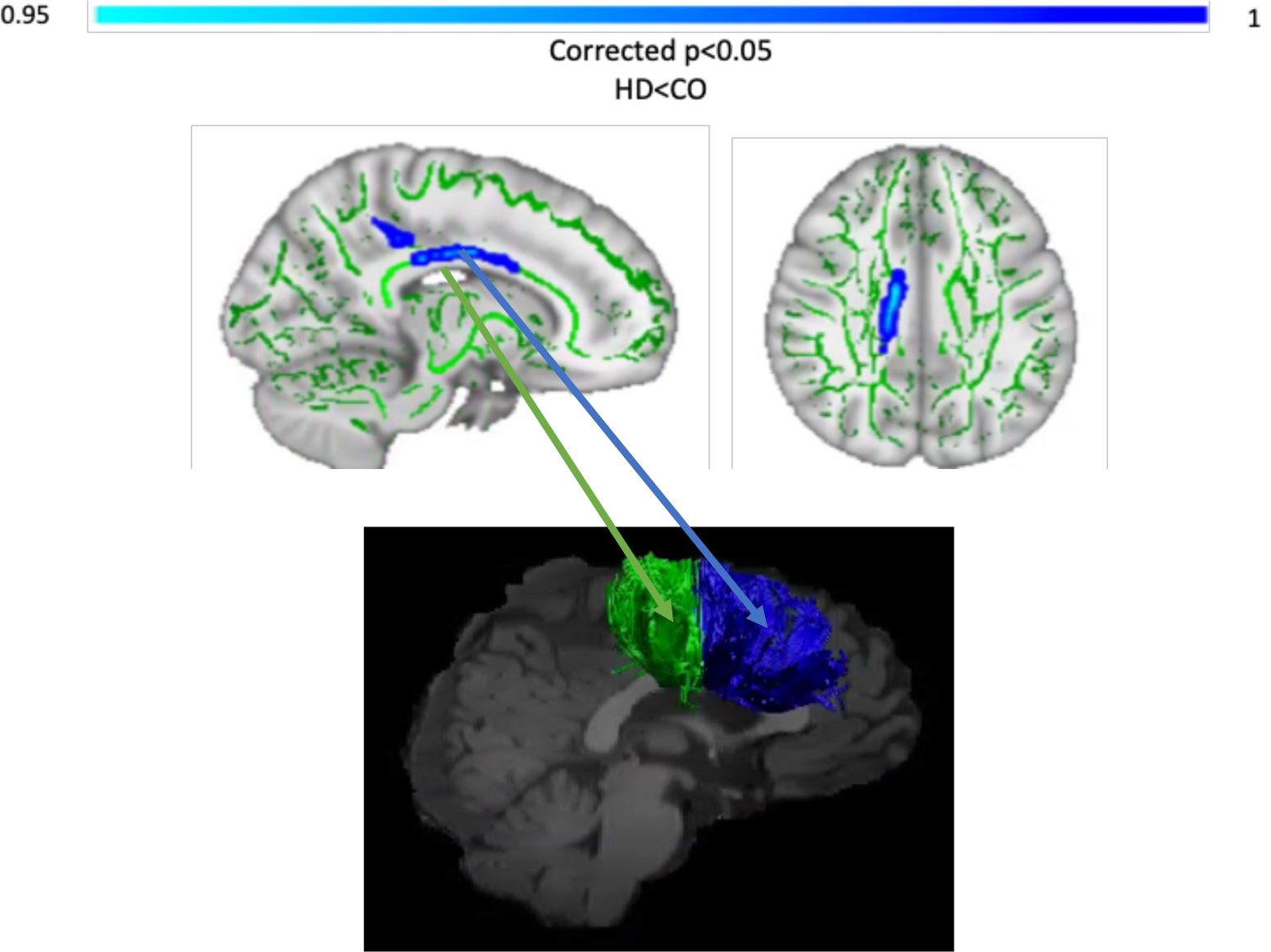
TBSS analysis of baseline MPF values. **(A)**. Light blue areas show a significant reduction of MPF in patients with HD compared to controls (p < 0.05, FWE corrected). The midbody of the CC was mostly found to be affected, which carries connections to the premotor, supplementary motor and motor areas of the brain. **Tracts showing significantly greater MPF changes in HD patients post-training as compared to controls (B)**. Areas showing significant MPF reductions at baseline overlap with tracts showing significant changes post-training (i.e. CCII and CCIII).

## 5. Discussion

Based on evidence that myelin impairment contributes to WM damage in HD (Bartzokis et al., 2007), and the suggestion that myelin plasticity underlies the learning of new motor skills (Lakhani et al., 2016; Scholz et al, 2009), the present study explored whether two months of drumming training would result in changes in WM microstructure in HD patients. Specifically, we expected to detect changes in MPF, as marker of WM myelin plasticity, in HD patients relative to healthy controls.

Firstly, we demonstrated a behavioural effect of the training by showing a significant improvement in drumming performance in patients (easy test pattern) and controls (medium test pattern). We did not detect any group differences in training-associated changes in the diffusion based indices of FA, RD and Fr. However, as hypothesised, we found a group difference in training-induced changes in the MPF PCA component. Specifically, HD patients showed significantly higher increases in MPF relative to controls. Furthermore, through exploratory post-hoc investigations, we detected significantly higher training-induced MPF changes within the CCII, CCIII and the right SMA-putamen pathway between patients and controls. Additionally, TBSS analysis of baseline differences in MPF suggested partial overlap of WM areas showing significant MPF reductions at baseline with areas showing changes post-training (i.e. CCII and CCIII).

MPF can be affected by inflammation (Henkelman, Stanisz, & Graham, 2001) and in advanced HD it is likely that inflammation goes hand-in-hand with myelin breakdown (Rocha et al., 2016). However, a recent CSF biomarker study found no evidence of neuro-inflammation in early-manifest HD (Vinther-Jensen et al., 2016). Furthermore, although recent evidence shows that this measure may be inconsistent when investigated in relatively small WM areas, presumably because of the effect of spatial heterogeneity in myelin thickness (Wang et al., 2020), we think that assessing within-subjects changes in MPF might have helped to address this limitation. Therefore, though preliminary and based on a small sample size, our findings suggest that two months of drumming and rhythm exercises may result in myelin remodelling in patients with early HD.

It is plausible that this group difference arose due to WM microstructural differences between patients and controls before the training. Accordingly, the HD group showed a significantly lower baseline MPF, consistent with lower myelin content (Bartzokis, 2007). Furthermore, previous studies have reported that training-associated percentage changes in MRI measures tend to be higher in patients than in healthy subjects (Caeyenberghs et al., 2018). One possibility is that in the healthy brain, neural networks may be optimally myelinated and further increasing myelin may not improve performance (Chomiak & Hu, 2009; Kaller et al., 2017; Rushton, 1951). Disentangling the impact of prior WM microstructural differences on microstructural plasticity during learning is beyond the scope of the current work.

Notably, the behavioral effect of drumming training and cognition differed between patients and controls. Patients improved in the easy drumming test pattern, and control improved in the medium test pattern. Furthermore, consistent with evidence from our pilot study (Metzler-Baddeley et al., 2014), patients showed increases in the executive function components whilst control participants did not show improvements in their cognition. Therefore, inter-group differences in microstructural changes might not only be due to baseline WM microstructural differences, but also to a different behavioral effect of the task between HD subjects and controls. For instance, control participants performed close to ceiling in the easy test pattern, and as the training was tailored to patients’ needs, some of the earlier practice sessions may not have optimally challenged them. The fact that the training seemed more taxing for patients than controls may also explain why improvements in executive functions and changes in MPF were only observed for the patients.

A critical question relevant to all training studies concerns the functional significance of any observed neural changes. While we expected significant relationships between changes in MRI measures and changes in drumming proficiency or performance in cognitive tests, no significant correlations were found. This might have been due to non-specific training-related neural responses. Specifically, while the training exercise might have triggered changes in brain structure, training-induced changes may not necessarily co-vary with improvements in performance. Alternatively, it might be that our study was insufficiently powered to detect brain-function correlations. The minimum sample size required to detect a correlation was calculated to be 64 people (α = 0.05; 80% power; medium effect size; GPower 3 software). Therefore, our results need replication in larger samples. Furthermore, lack of correlation between structural and functional changes after training has been reported in other studies (including well-powered studies) and may suggest that these processes follow different time courses and may occur in different brain regions (Valkanova et al., 2014).

It is important to note that our study did not include a non-intervention patient group. Within the 12 month time period of this study it was not possible to recruit a sufficiently large number of well-matched patient controls. Therefore, we cannot disentangle the effects of the training on WM microstructure from HD-associated pathological changes. However, given that HD is a progressive neurodegenerative disease associated with demyelination (Bartzokis et al., 2007), it is unlikely that increases in MPF observed in the patient group were due to the disease itself. Finally, while the majority of training studies assess brain structural changes between baseline and post-training (Caeyenberghs et al., 2018), presumably on account of cost and participant compliance, we suggest that acquiring intermittent scans during the training period could have helped to better capture and understand changes in WM microstructure observed in this study. Accordingly, future studies and more advanced statistical analyses might be able to give greater insights into the complex nonlinear relationships between structural changes and behaviour (Thomas & Baker, 2013).

To conclude, we have demonstrated that two months of drumming and rhythm exercises result in a significantly greater change in a proxy MRI measure of myelin in patients with HD relative to healthy controls. Whilst the current results require replication in a larger patient group with an appropriately matched patient control group, they suggest that behavioural stimulation may result in neural benefits in HD that could be exploited for future therapeutics aiming to delay disease progression.

## List of abbreviations

ANOVA: analysis of variance
CC: corpus callosum
CHARMED: Composite Hindered and Restricted Model of Diffusion
CI: confidence intervals
DT MRI: diffusion tensor magnetic resonance imaging
DTI: diffusion tensor imaging
EPI: echo planar imaging
FA: fractional anisotropy
FAS: functional assessment score
FDR: false discovery rate
fODF: fiber orientation density function
FoV: field of view
FR: restricted fraction
FSPGR: fast spoiled gradient echo
FWE: family-wise error
GM: grey matter
HD: Huntington’s disease
MPF: macromolecular proton fraction
MRI: magnetic resonance imaging
MT: magnetization transfer
MT-w: MT-weighted
PCA: principal component analysis
qMT: quantitative magnetization transfer
RD: radial diffusivity
ROI: region of interest
SD: standard deviation
SE: spin-echo
SMA: supplementary motor area
SPGR: spoiled gradient recalled-echo
T1-w: T1-weighted
TBSS: tract-based spatial statistics
TE: echo time
TMS: total motor score
TR: repetition time
UHDRS: Unified Huntington’s disease rating scale
WM: white matter

## Declarations

### Ethics approval and consent to participate

The study was approved by the local National Health Service (NHS) Research Ethics Committee (Wales REC 1 13/WA/0326) and all participants provided written informed consent.

### Availability of data and materials

The datasets analysed during the current study are available from the corresponding author on reasonable request.

## Funding & Acknowledgements

The present research was funded by a Wellcome Trust Institutional Strategic Support Fund Award (ref: 506408) to CMB, AR, and DKJ and a Wellcome Trust PhD studentship to CC (ref: 204005/Z/16/Z); DKJ is supported by a New Investigator Award from the Wellcome Trust (ref: 096646/Z/11/Z). We would like to thank Candace Ferman and Louise Gethin for collating the patients’ clinical details and Jilu Mole for assistance with data analysis. This manuscript has been released as a pre-print at bioRxiv: https://biorxiv.org/cgi/content/short/2019.12.24.887406v1

